# Postponing production exponentially enhances the molecular memory of a stochastic switch

**DOI:** 10.1101/2020.06.19.160754

**Authors:** Pavol Bokes

## Abstract

Delayed production can substantially alter the qualitative behaviour of feedback systems. Motivated by stochastic mechanisms in gene expression, we consider a protein molecule which is produced in randomly timed bursts, requires an exponentially distributed time to activate, and then partakes in positive regulation of its burst frequency. Asymptotically analysing the underlying master equation in the large-delay regime, we provide tractable approximations to time-dependent probability distributions of molecular copy numbers. Importantly, the presented analysis demonstrates that positive feedback systems with large production delays can constitute a stable toggle switch even if they operate with low copy numbers of active molecules.

## 1 Introduction

Protein products of a given gene can be present in single cells at low copy numbers, fluctuating in time due to random occurrences of production and degradation events [1–3]. The resulting protein probability distributions exhibit large ratios of variance to the mean, especially if the protein is produced in bursts [4–8]. Understanding how cells can maintain predictable behaviour in the presence of gene-expression noise poses an important challenge to mathematical and systems biology.

The completion of a functioning protein molecule involves a succession of processes such as transcriptional and translational elongation [9, 10], post-translational modification [11], and splicing and export to cytoplasm in case of eukaryotes [12–14]; any of these effectively postpones the production event. Delayed production can dramatically alter the protein dynamics, especially if the protein autoregulates its own production [15–17]. While delays in negative feedback systems can lead to destabilisation and noise amplification [18–20], delays have been reported to stabilise the behaviour of and reduce noise in positive feedback systems [21–24]. Long distributed delays buffer production bursts and reduce the large protein variance-to-mean ratios to the Poissonian value of one [25–28]. If a large exponentially distributed delay is added to a highly cooperative positive feedback system, the protein distribution is a mixture of multiple Poissonian modes [29]. In this paper, we extend the steady-state result of [29] and characterise the temporal evolution of the Poissonian modes.

The key insight of [29], on which we expand in this work, is that a stochastic autoregulatory circuit extended by a large delay exhibits a near-deterministic behaviour of the inactive protein (the production of which has been initiated but not yet completed). A classical means of eliminating noise in a stochastic model is to increase its system size/volume [30]; the near-deterministic behaviour in large-volume systems can be described using the linear noise approximation (LNA) [31–35] and the Wentzel–Kramers–Brillouin (WKB) approximation [36–40]. These two approaches are complementary: the LNA applies on finite temporal domains [41]; the WKB approximation covers slow metastable dynamics such as transitions between deterministically stable steady states or to a fixation/extinction point [42]. The LNA and WKB methodologies have been extended to large-volume systems which additionally include a fast-equilibrating low-copy number species [43–47]. While the assumptions of high abundance of slow species and fast equilibration of scarce species are both required for a deterministic limit, other studies considered the high-abundance limit without eliminating the fast species [48–51], or the fast equilibration limit without assuming high abundance in the slow species [52–54]. The lengthening of production delay in [29] has the combined effect of making the inactive protein highly abundant and the scarce active protein (relatively) fast; this renders large-delay systems tractable to LNA and WKB methods.

The outline of the paper is as follows. Section 2 introduces the delayed feedback model and its master equation. Sections 3, 4, and 5 develop the LNA, the WKB-based mixture approximation, and an initial-transient approximation to the solution of the master equation, respectively. Section 6 concludes the paper with a discussion.

## 2 Master equation

The numbers of inactive (*X*) and active (*s*) proteins are modelled by a bi-variate Markov chain in continuous time with discrete state space {0,1,…} × {0,1,…, *s*_max_}. Here we are making a modelling assumption that the number of active proteins *s* stays below an upper bound *s*_max_; this will enable us to use finite-dimensional techniques in our analysis later. The assumption is not restrictive, as a wide class of unbounded systems can be arbitrarily closely approximated by truncated systems [55–57].

The joint probability mass function (PMF) *P*(*X, s, t*) of *X* and *s* satisfies the Kolmogorov forward equation [58]

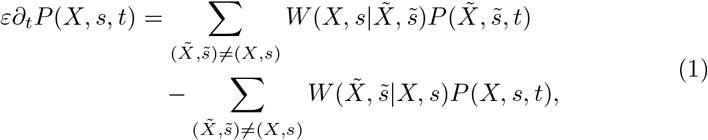

where 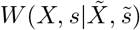 gives the transition rates in the Markov chain; they are of the following types:

### Production

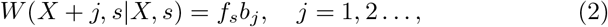

in which *b_j_* ≥ 0, where 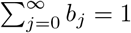, gives the probability of a production burst of size *j* = 0, 1,…; the dependence of the production rate *f_s_* on the active protein *s* implements the delayed feedback.

### Activation

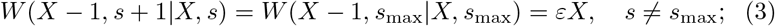

the boundary modification of the activation transition guarantees that the number *s* of active proteins never exceeds the upper bound *s*_max_. The activation rate constant *ε* (which is the reciprocal of the mean delay) is assumed to be small.

### Decay

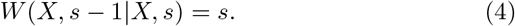

The Kolmogorov forward equation (1) with transitions (2)–(4) reads

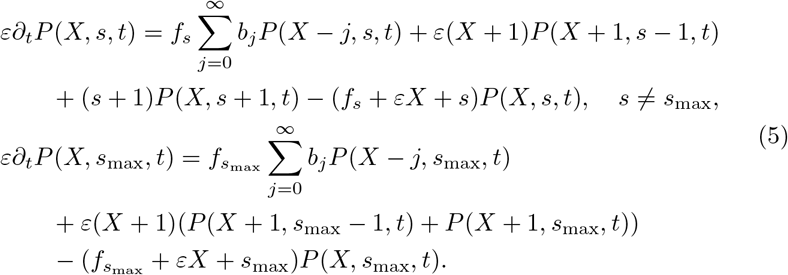

The probability of having a negative amount of *X* or *s* is tacitly understood to be equal to zero in (5) and equations that follow. The aim of the paper is to provide asymptotic approximations as *ε* → 0 for time-dependent solutions *P*(*X, s, t*) to (5).

Expressing the shifts in the variable *X* using the exponential of the differential operator [36]

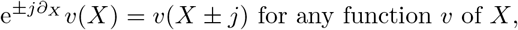

and defining the moment-generating function (MGF) of production burst size by

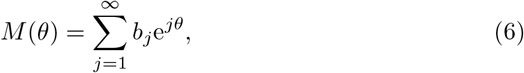

we rewrite (5) as

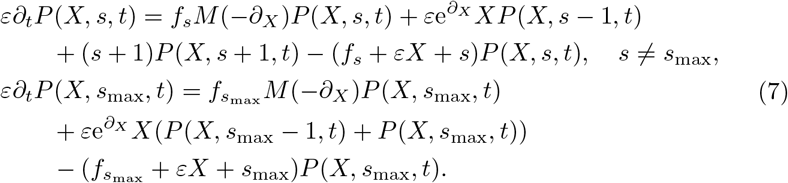

Since the inactive protein is produced at an *O*(1) rate but is removed (by being activated) at an *O*(*ε*) rate, its copy number *X* will accumulate to *O*(1/*ε*)-large levels. In order to bring the inactive protein measurements back on the *O*(1) scale, we introduce new variables

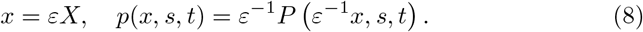

We refer to *x*, which becomes a continuous quantity in the limit of *ε* → 0, as the concentration of the inactive protein. The *ε*^−1^ prefactor is included in the second equality of (8) so that *p*(*x, s, t*) represents the probability density function (PDF) of *x*, rather than its PMF. Inserting (8) into (7) gives

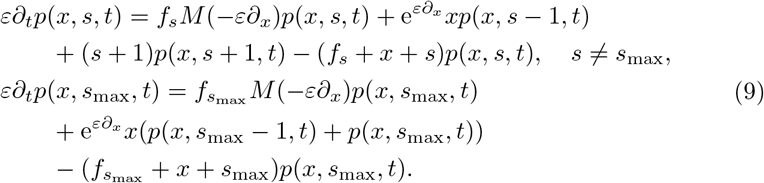

Equation (9) can be vectorised into

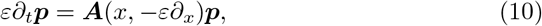

where

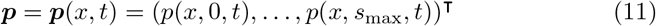

and

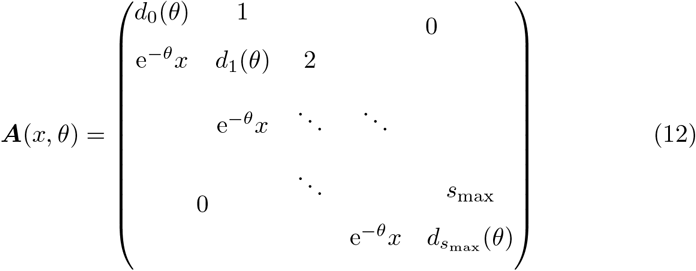

is a tridiagonal matrix with main diagonal elements defined by

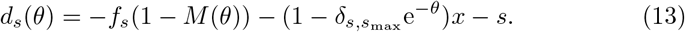

The insertion in (10) of the differential operator −*ε∂_x_* in place of the parameter *θ* of the matrix (12)–(13) has one important subtlety: the composition of operators, unlike multiplication, is not commutative; in particular, *e^−θ^x ≠ xe^−θ^* in (12)–(13). Equation (10), being the focal point of much of the ensuing analysis, will be referred to as the master equation.

## 3 Linear noise approximation

We express the inactive protein concentration as the sum of a deterministic function of time and a small random correction

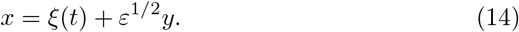

Denote by Π(*y, s, t*) the joint probability distribution of the inactive protein fluctuation *y* and the active protein copy number *s*, and let

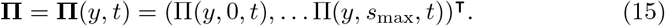

By the transformation rule, the distribution of *y* and that of *x* are related via

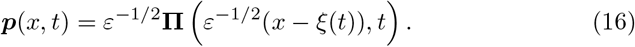

The inclusion of the prefactor *ε*^−1/2^ in (16) ensures that the distribution remains normalised after transformation. In the expansion procedure that follows, we determine *ξ*(*t*) and, to the leading order, **Π**(*y,t*); equation (16) then reconstitutes the approximation in terms of the original joint distribution ***p***(*x, t*).

### 3.1 Expansion

Inserting the transformation (14) and (16) into the master equation (10) and multiplying by *ε*^1/2^ yield

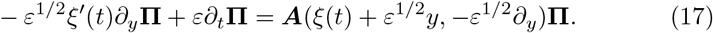

Taylor-expanding the right-hand side of (17),

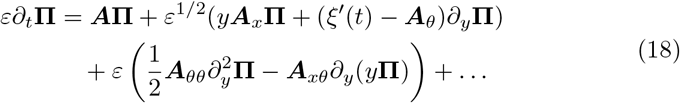

The matrix ***A*** and its derivatives ***A**_x_, **A**_θ_, **A**_θθ_, and **A**_xθ_* are understood to be evaluated at *x = ξ*(*t*), *θ* = 0 in (18) and equations that follow; note that, since the matrix elements (12)–(13) depend linearly on *x*, we have ***A**_xx_* = 0.

We look for solutions to (18) in the form of a regular power series in *ε*^1/2^, i.e.

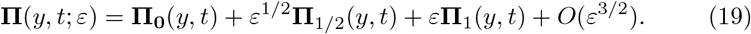

Inserting (19) into (18) and collecting terms of same order yield

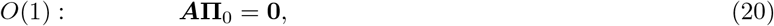

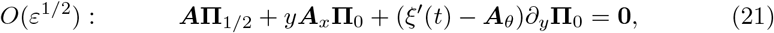

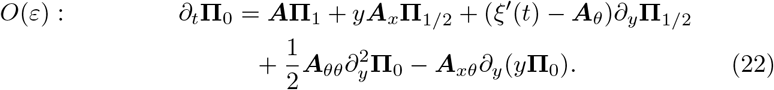

In the next subsections we go through the consequences of equations (20)–(22), one by one.

### 3.2 The active protein probability mass function

By (20), the leading-order approximation **Π**_0_ = **Π**_0_(*y, t*) to the joint protein distribution belongs to the nullspace of the matrix ***A = A***(*ξ*(*t*), 0). Substituting *θ* = 0 into (12)–(13), whilst noting that *M*(0) = 1, one realises that the matrix ***A***(*x*, 0) is the Markovian generator matrix of a one-dimensional truncated birth–death process (Figure 1). The nullspace of ***A***(*x*, 0) consists of scalar multiples of its stationary distribution ***ρ***(*x*) = (*ρ*_0_(*x*),…, *ρ*_*s*_max__(*x*))^**⊤**^, which is the truncated Poisson distribution

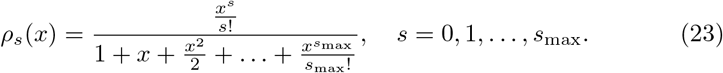

The leading-order approximation to the joint protein distribution therefore satisfies

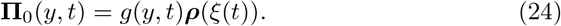

**Figure 1:**
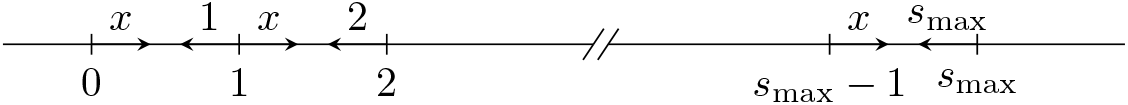
The one-dimensional truncated birth–death process with constant birth rate *x* and linear death rate *s*. The discrete states correspond to the admissible copy numbers *s* of the active protein, births represent activation (here from an unchanging pool of inactive proteins), and deaths correspond to decay.

By (24), the inactive protein concentration fluctuation *y* and the active protein copy number *s* are at any time *t* statistically independent; the probability mass function of the latter is given by (23); the probability density function of the former is given by the function *g*(*y, t*), which will be determined in Subsection 3.4. Before doing that, let us identify in the next subsection the deterministic motion ξ(*t*) of the inactive protein concentration.

### 3.3 The deterministic motion of the inactive protein

Inserting (24) into (21), one finds

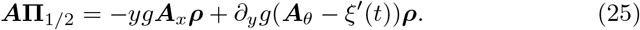

Multiplying (25) from the left by the (*s*_max_ + 1)-dimensional row vector **1^⊤^** = (1,…, 1) of ones, and noting that **1^⊤^*A* = 1^⊤^*A***_*x*_ = **0** and **1^⊤^*ρ* = 1**, we realise that in order that (25) be solvable in **Π**_1/2_, it is necessary that *∂_y_g*(**1^⊤^ *A***_*θ*_***ρ***−*ξ′*(*t*)) = 0 holds; and, as we do not allow *g* to be constant in *y*, this means that

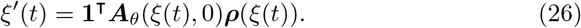

Next, we simplify the right-hand side of the ordinary differential equation (26). Differentiating the elements of the matrix (12) with respect to *θ*, then taking *θ* = 0 whilst noting that *M′*(0) = 〈*B*〉, we find, after a little algebra, that

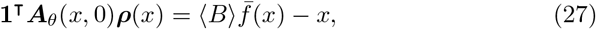

where

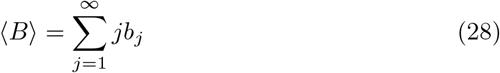

gives the mean burst size and

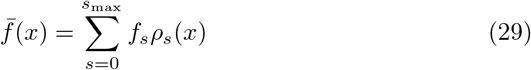

gives the effective production rate. Formula (29) calculates the average of the instantaneous production rate with respect to the Poisson distribution (23) of the active protein copy number.

Inserting (27) into (26), we find

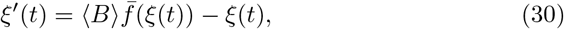

meaning that the deterministic component *ξ*(*t*) of the inactive protein concentration increases per unit time by the effective production rate and descreases by the linear activation rate.

### 3.4 The inactive protein fluctuation

Inserting (24) and (26) into (22) and multiplying from the left by **1^⊤^** yield

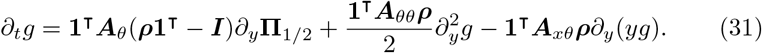

In order to close (31), we need to express **Π**_1/2_ in terms of the unknown g. The right-hand side of (25) is a linear combination whose coefficients depend on y but the vectors do not. We write the solution to (25) as

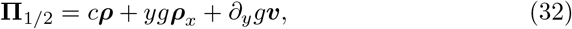

where *c* is a scalar (possibly dependent on *y* and *t*), ***ρ*** is a homogeneous solution, and ***ρ**_x_* and ***υ*** solve the y-independent problems

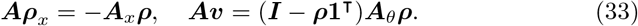

That ***ρ**_x_* is indeed a solution to the first of the inhomogeneous problems (33) can be established by differentiating ***Aρ* = 0**.

Inserting (32) into (31) whilst noting that **1^⊤^*ρ***_*x*_ = 1_*x*_ = 0 yields a linear time-dependent Fokker–Planck equation

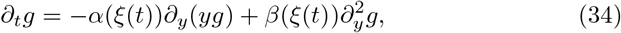

where

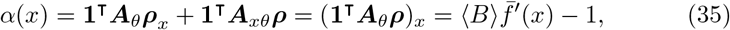

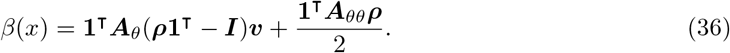

The drift term *α*(*x*) is the linearisation of the deterministic motion (30). While no apparent simplification is available for the first term in (36), the second term satisfies 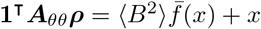.

Assuming without loss of generality that at the initial time *t* = 0 there is no deviation of the stochastic process from the initial condition *ξ*(0) = *x*_init_ of the deterministic equation (30), we solve the Fokker–Planck equation (34) subject to the delta-function initial condition

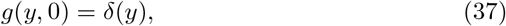

obtaining a Gaussian solution [32]

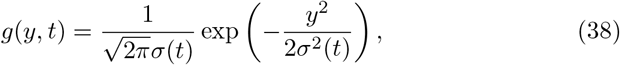

where the variance *σ*^2^(*t*) satisfies an inhomogeneous linear equation of the first order with variable coefficients

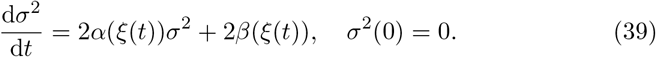

Equation (39) is in related contexts referred to as the fluctuation–dissipation relation [59]

### 3.5 The LNA and its small-*t* and large-*t* behaviour

Combining (16), (19), (24), and (38), we arrive at the final form of the linear noise approximation (LNA)

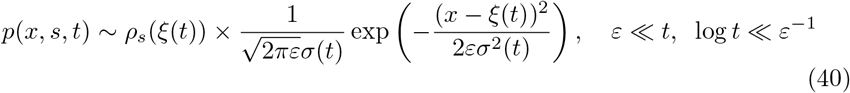

for the joint distribution of the concentration *x* of the inactive and the copy number *s* of the active protein. The LNA (40) implies that: *s* follows the truncated Poisson distribution (23); *x* follows the Gaussian distribution with mean equal to the deterministic solution *ξ*(*t*) to (30) and a small variance equal to *εσ*^2^(*t*).

Although the analysis in the current section implicitly assumes *t* = *O*(1), the validity of (40) extends to all times except for those that are *O*(*ε*) small or whose logarithms are *O*(1/*ε*) large (i.e. exponentially large times). For these two time domains different approximations will be developed in Sections 4 and 5. In preparation for these developments, note that the asymptotic matching principle [60] implies that the terminal behaviour of an early-time approximation has to coincide with the *t* ≪ 1 behaviour of (40), and the *t* ≫ 1 behaviour of (40) with the initial behaviour of a large-time approximation. The former is

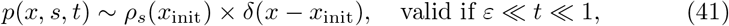

as can be seen easily by taking the limit of *t* → 0 of (40).

The *t* → ∞ limit of (40) deserves a more detailed treatment especially if the deterministic motion (30) is bistable (which we assume to be the case henceforth). Bistability can occur if there is a positive feedback in protein production, and the effective production rate is sufficiently sigmoid (Figure 2, left). The deterministic model (30) then possesses three steady states *x*_−_ < *x*_0_ < *x*_+_, of which *x*_−_ and *x*_+_ are stable and *x*_0_ is unstable (Figure 2, left, diamonds). Depending on which side of the unstable fixed point *x*_0_ the initial protein concentration *ξ*(0) = *x*_init_ resides, the deterministic solution (30) and the variance coefficient (39) assume different large-time limits

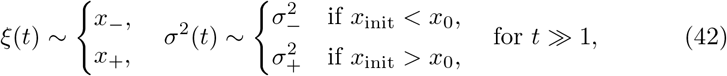

in which

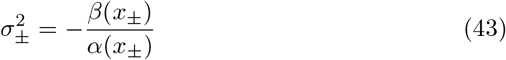

are the two possible large-time limits of (39). With *β*(*x*) being positive and *α*(*x*) negative at the stable steady states *x = x*_±_, the variances (43) are positive as should be. Note that we discarded from our considerations the non-generic possibility *x*_init_ = *x*_0_.

**Figure 2:**
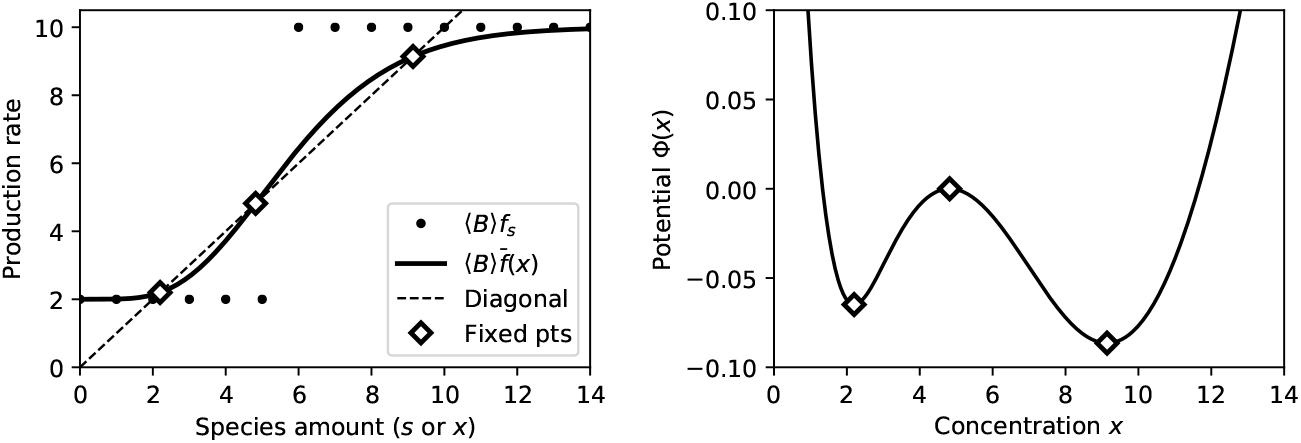
*Left:* The instantaneous production rate 〈*B*〉*f_s_* (dots) and the Poisson-averaged production rate 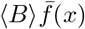 (full line). As a result of the averaging (29), the sharp features of the former are smoothed out in the latter. Fixed points *x*_−_ < *x*_0_ < *x*_+_ (diamonds) of the deterministic model (30) occur where 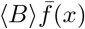 intersects the diagonal (dashed line). *Right:* The WKB potential possesses local extrema where (30) has fixed points. *Parameters for both panels:* the burst rate is *f_s_* = 0.5 for *s* ≤ 5; *f_s_* = 2.5 for *s* ≥ 6; the burst size is *B* = 4 (*b_j_ = δ*_*j*,4_).

Inserting (42) into (40) yields

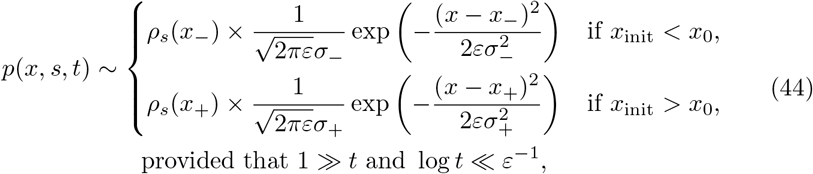

which can be expressed in terms of the Heaviside function in a single formula

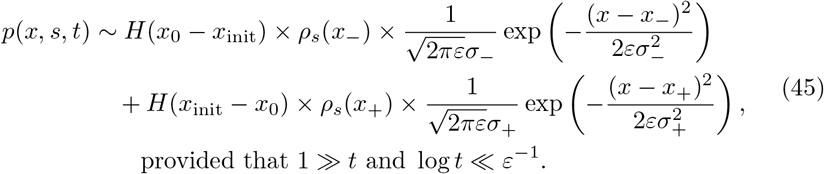

The resulting approximation (45) is a mixture of two Poisson/Gaussian modes located at the stable fixed points of the deterministic rate equation (30). The weights in (45) are binary: the whole probability mass is given to one or the other peak depending on the initial condition. On the exponentially large timescale, however, the weights of the peaks become probabilistic and slowly evolve in time, as described in the next section.

## 4 Mixture approximation

The large-timescale approximation to the joint probability distribution is obtained by replacing the binary weights in (45) by time-dependent ones,

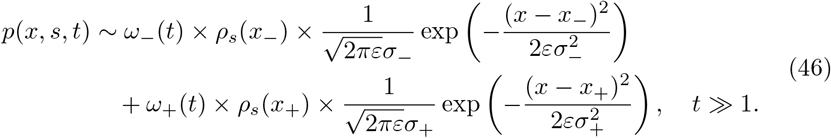

The weights are set to evolve according to a random telegraph process

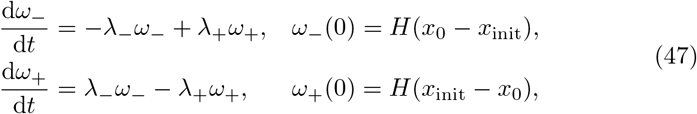

in which λ_±_ are the metastable transition rates between the modes of the distributions. The weight values are initiated in (47) to the binary values that were identified by the large-*t* analysis of the LNA.

The explicit solution to the random-telegraph equations (47) is given by

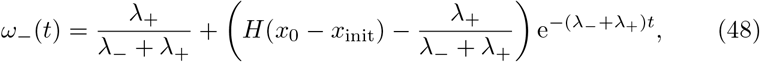

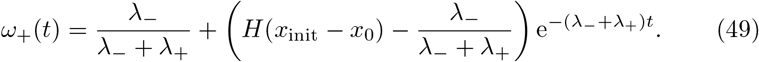

With (46) and (48)–(49) at hand, it remains to determine the metastable transition rates λ_±_ to complete the large-time approximation to the joint protein distribution. This is done in the subsections that follow; at the present moment, we point out that the rates will be shown to be exponentially small,

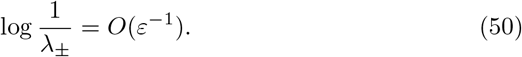

It follows that unless time *t* is exponentially large, the product (λ_+_ + λ_−_)*t* in the exponential in (48)–(49) is negligible, so that

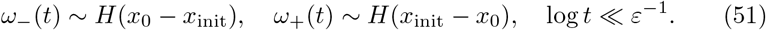

Note that inserting (51) into (46) recovers (45). Thus, provided that time is large but not exponentially large, the mixture distribution (46) with randomtelegraph weights and the LNA approximation (40) do indeed overlap.

The LNA (40) and the mixture approximation (46) to the probability distribution *p*(*x, s, t*) are easily recast, by means of the transformation (8), into approximations for the joint probability mass function *P*(*X, s, t*) of the copy numbers *x* and *s*. In order to validate the approximations, we also calculate the PMF numerically. This requires restricting the model in *x* like we did in *s*, i.e. to a finite state space {0,1,…, *X*_max_} × {0, 1,…, *s*_max_}. For the finite process, the bursty increases (2) apply only if the burst size satisfies *j* < *X*_max_ − *X*, with larger bursts being redirected to the boundary as per 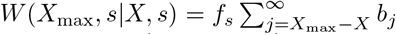. The Kolmogorov forward equation (1) is then a finite (albeit large) linear system of differential equations, and is solved by numerical integration.

**Figure 3:**
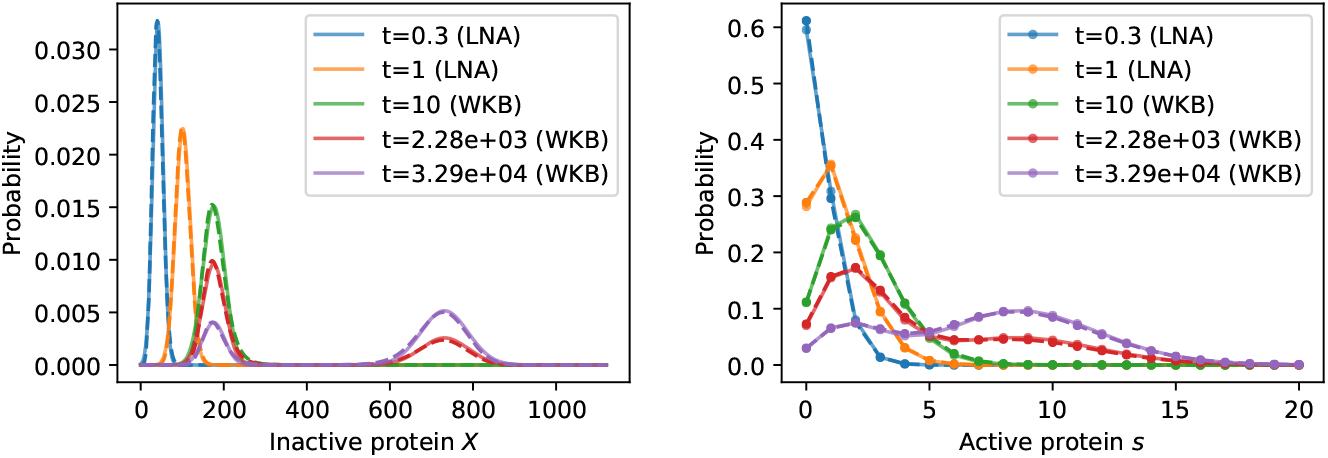
Numeric (*dashed line*) and asymptotic (*full line*) approximations to the PMFs of the inactive (*left panel*) and active (*right panel*) protein. The markers indicate the discrete values of the active protein distribution. The LNA (40) is used for early, and the WKB-based approximation (46) for the later timepoints; the last two time-points are set to log(2)/(λ_−_ + λ_+_) and 10/(λ_−_ + λ_+_), where λ_±_ are the metastable transition rates (63). The model parameters are *f_s_* = 0.5 for *s* ≤ 5; *f_s_* = 2.5 for *s* ≥ 6; *B* = 4; *ε* = 0.0125; *s*_max_ = 20. The initial values are *X*_init_ = *s*_init_ = 0. The upper bound for *x* required for the numerical solution is *X*_max_ = 1121.

Figure 3 shows the evolution of the marginals of a solution which is initialised to zero copy numbers, i.e. *P*(*X, s*, 0) = *δ*_*X*,0_*δ*_*s*,0_. The zero initial state lies in the region of attraction of the lower stable fixed point *x*_−_. The early dynamics is characterised by a steady shift of the distributions to the right until they are located at *x*_−_. This initial dynamics is closely approximated by the LNA (40) (Figure 3, first three timepoints). At much greater times, the weight of the mode around *x*_−_ begins to decrease, and a new mode around the upper stable fixed point *x*_+_ arises. This later dynamics is closely approximated by the mixture approximation (46) (Figure 3, last three timepoints); for reasons that will become apparent in the next section we refer to the mixture as the WKB-based approximation. At the final timepoint, a steady state is reached, which is characterised by a balance of probability transfer between the two modes.

### 4.1 WKB approximation

The determination of λ_±_ rests on a uniformly valid approximation to the stationary distribution that has been provided in an earlier work [29] in the WKB

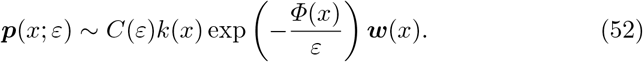

For a detailed exposition how the individual terms in (52) can be calculated we refer the reader to the original paper. Here we merely summarise the most important properties of these terms. The function *Φ*(*x*), which is referred to as the potential, plays a key role in the approximation. Small differences in the potential are exponentially exacerbated in (52); it follows that the main contribution toward the total probability mass comes from around the local minima of the potential. Importantly, [29] showed that *Φ*(*x*) has local minima at the stable fixed points *x*_±_ of the differential equation (30), separated by a local maximum at the unstable fixed point *x*_0_ (Figure 2, right panel). The vector ***w***(*x*) > 0 in (52) satisfies **1^⊤^*w***(*x*) = 1 and its components provide an approximation to the stationary distribution of the active protein copy number conditioned on a given concentration *x* of the inactive protein. The function *k*(*x*) > 0 is a prefactor, and the coefficient *C*(*ε*) > 0 is a normalisation constant.

Assume for concreteness that the protein concentration is initialised at *t* = 0 to a value *x*_init_ < *x*_0_. After the elapse of an initial *O*(1) temporal transient, the concentration *x* equilibrates to a quasi-stationary distribution located around the lower fixed point *x*_−_. The quasi-stationary distribution coincides with the stationary distribution (52) for *x < x*_0_, and is equal to zero for *x > x*_0_; the normalisation constant *C*(*ε*) is therefore calculated by integrating (52) over 0 < *x* < *x*_0_ only, i.e.

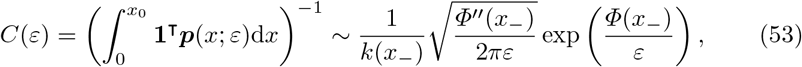

in which the approximation is due to the Laplace method. The truncation of (52) to zero for *x > x*_0_ introduces a discontinuity at *x*_0_ in the quasi-steadystate distribution. The discontinuity can be resolved by constructing in a small neighbourhood of *x*_0_ an inner approximation to the quasi-stationary distribution that asymptotically matches (52) to its left and zero to its right. Importantly, the construction of the inner solution and the matching procedure will help quantify the rate of probability leakage λ_−_ across the potential barrier *x*_0_. This is done in the next two subsections.

The quasi-stationary distribution can be alternatively approximated by the large-*t* limit of the LNA (44). Indeed, comparing the behaviour of (44) and (52) around the stable fixed points, one derives consistency relations

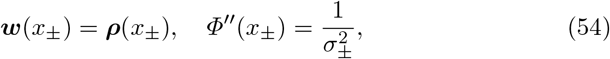

which have previously been obtained by [29] by other means. The first equality of (54) means that the WKB and LNA approaches consistently predict the same Poisson distributions of active protein if conditioned on a fixed-point concentration of the inactive protein; the second equality of (54) says that the linearised noise around a fixed point increases as the WKB potential well flattens. While the LNA and the WKB approximation are both appropriate around the fixed points, only the WKB approximation provides an adequate approximation at the tails of the distribution away from the fixed points. Since the exchange of probability mass between the fixed points occur through the tails, it is necessary to use the uniformly valid WKB approximation to obtain the correct rate of leakage. Nevertheless, the LNA methodology, aside from providing time-dependent approximations for *t = O*(1), plays a key role in the construction of the inner approximation to the quasi-stationary distribution, as explained below.

### 4.2 Inner approximation

Applying the transformation (14) with the deterministic motion set constantly to the unstable fixed point *ξ*(*t*) ≡ *x*_0_, and carrying through with the expansion procedure, we approximate via (16), (19), and (24) the quasi-stationary distribution by

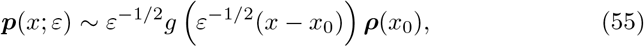

where *g = g*(*y*) is a solution to the time-independent version of the Fokker–Planck equation (34),

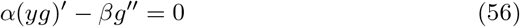

with *α = α*(*x*_0_) and *β* = *β*(*x*_0_) given by (35)–(36). Reducing the order of (56) gives

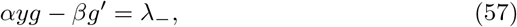

where λ_−_ is a constant of integration. Solving (57) in *g* subject to the terminal condition *g*(∞) = 0 yields

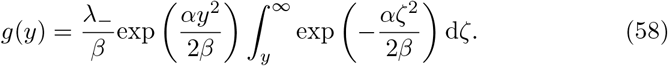

The integration constant λ_−_ is identified with the metastable transition rate as it gives the (drift-diffusion) probability current — the amount of probability that is being transferred per unit time through the unstable fixed point in the direction of increasing *x*. The value of the probability current is determined in the next subsection by an asymptotic matching of (58) to the WKB formula for the quasi-stationary distribution.

### 4.3 Asymptotic matching

The WKB approximation (52) and (53) to the quasi-stationary distribution applies for values of *x* that lie to the left of the unstable fixed point *x*_0_. On the other hand, the LNA approximation (55) and (58) applies for *x* in the vicinity of *x*_0_. Asymptotic matching principle states that the two approximations should be equivalent in an overlap region, which is here given by

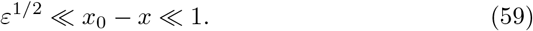

The LNA approximation (55) and (58) simplifies in the overlap region (59) to

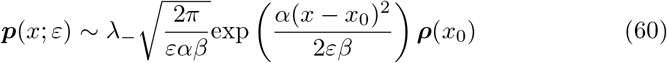

and the WKB approximation (52) and (53) to

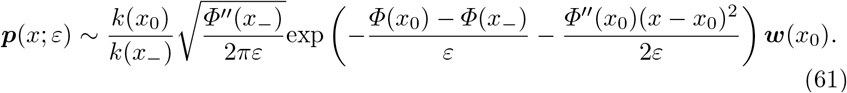

Comparing (60) and (61) yields

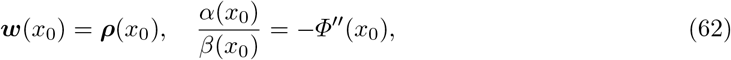

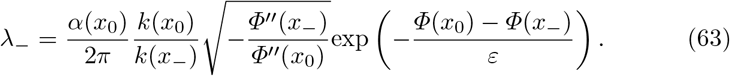

Since *Φ*(*x*_0_) > *Φ*(*x*_−_), the current (63) is exponentially small as *ε* → 0 as advertised in (50). The WKB terms *k*(*x*_−_), *k*(*x*_0_), and *Φ*(*x*_0_) − *Φ*(*x*_+_) can be numerically determined as described by [29]. The formula for λ_+_ is obtained by replacing the symbol ‘−’ by ‘+’ wherever it appears in formula (63).

## 5 Early-time approximation

For completeness, we provide an approximation to the joint protein distribution that applies during the small initial transient *t = O*(*ε*). First, we introduce the fast time variable by

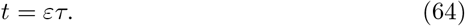

Inserting (64) into the master equation (10) yields

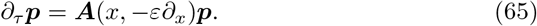

Taking *ε* → 0 in (65), and solving the limiting equation yields an approximation

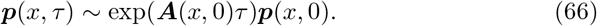

Expressing (66) in terms of the original time variable for deterministic initial conditions, we arrive at

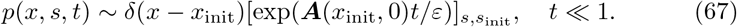

The exponential of the birth–death process’s (Figure 1) Markovian generator matrix in (67) can for *ε* ≪ *t* be approximated by the its stationary distribution *ρ_s_*(*x*_init_); making this approximation recovers (41). Thus, provided that time is small but sufficiently larger than *ε*, the birth–death process dynamics of (67) and the LNA approximation (40) do indeed overlap.

## 6 Discussion

The paper describes the stochastic dynamics of a bursty protein with delayed feedback in the large-delay regime. A key observation is that long delays lead to a large build-up of protein molecules which have already been produced but not yet activated. The reservoir of inactive protein molecules confers stability to random perturbations on the inactive protein. Indeed, in the limit of very large delays, the inactive protein evolves deterministically according to an autonomous ordinary differential equation (30). Meanwhile, the number of active protein fluctuates fast, being “slaved” — borrowing the metaphor from [61] — by the inactive protein. Provided that the protein sustains its expression via a sufficiently cooperative positive feedback loop, the limiting differential equation possesses two stable fixed points, a lower one of basal expression and a higher one of up-regulated expression. Bistablility generated by a positive feedback can implement molecular memory in living cells and serve to maintain cell-fate decisions [62–65].

The delayed regulatory circuit is a multiscale problem with three distinct timescales: the fast timescale of the active protein turnover, the slow timescale of the inactive protein turnover, and the very slow timescale of metastable transitions between the fixed points of the deterministic motion. As the main mathematical result of the paper, we develop on each of the three timescales an approximation to the joint probability distribution of inactive and active protein abundances. On the fast timescale, the inactive protein stays constant, while the active protein equilibrates to the Poisson distribution (67). On the slow timescale, the inactive and active protein species follow respectively the Gaussian and the Poisson distributions (40) located around the deterministic motion. On the very slow timescale, the inactive protein distribution is a mixture of Gaussians, and the active protein distribution is a mixture of Poisson modes (46) located at the fixed points of the deterministic motion. The weights of the modes evolve in time according to a very slow two-state switch (a random telegraph process), the holding times of which exponentially lengthen as the delay increases. Therefore, even moderately large delays can implement an extremely stable switching mechanism that can function as a basic molecular memory unit in a living cell.

As part of our asymptotic analysis, we demonstrated that the successive timescales share a common overlap domain on which the respective approximations provide consistent results. On the one hand, the existence of the overlap demonstrates the completeness of our approximation scheme in the sense of covering the whole temporal frame; on the other hand, the requirement that the terminal behaviour of a preceding approximation is continued by the successive approximation determines the initial condition for the latter. Asymptotic matching is used not only in the temporal but also in the protein domain, where it serves to determine the rates of metastable escape from the basins of attraction of the deterministic fixed points. The procedure, which has previously been applied to large-volume [36, 37, 39] and fast-switching systems [44, 45], involves outer and inner approximations to the quasi-stationary protein distribution. The outer approximation, which applies within the basin, is calculated by the WKB method following [29]. The inner approximation, which is valid in a narrow neighbourhood of the unstable fixed point that separates the basins, is found as a constant-flux solution to a Fokker–Planck equation. The probability flux of the inner solution is determined from matching the inner to the outer approximation; the value of the flux provides the desired metastable rate of escape from the basin.

In summary, our study demonstrates that extending a noisy autoregulatory loop by a sufficiently large production delay can form a stable toggle switch. Using a matched asymptotic approach, our study quantifies the effect and connects the metastable switching to shorter-term dynamics. Finally, we expect that the phenomenon is present not only in case of a single gene, but also appears for mutually repressive genes, and other manifestations of positive feedback systems.

## Acknowledgement

The author thanks Abhyudai Singh, Pasquale Palumbo, and Alessandro Borri for discussions which influenced the paper.

